# Activity of oleo-gum resins of myrrha on the growth of Candidiasis

**DOI:** 10.1101/2023.12.22.572985

**Authors:** Amira H. Alabdalall

## Abstract

**Aims:** Medicinal plants have a long and significant history of being used for their healing properties. One famous example is Commiphora, which is mostly found in the southern part of Arabia. The objective of this research was to evaluate the effectiveness of a water-based extract obtained from two different varieties of myrrh in suppressing the proliferation of *Candida* spp. at different concentrations. The objective of this research is to determine the chemical composition of *Commiphora* and its capacity to prevent the development of *Candida* spp.

**Methodology and Results:** A plant specimen traditionally used in Saudi Arabia for its possible effectiveness against *Candida albicans* (ATCC 14053), *C. neoformans* (ATCC 66031), *Candida laurentii* (ATCC 18803), *Candida guilliermondii* (ATCC 6260), and *Candida tropicalis* (ATCC 66029). The results showed that the aqueous extract of both tested species of Myrrha (*Commiphora myrrha* and *C. molmol*) shown inhibitory effects on all tested isolates. Moreover, it was discovered that the inhibitory impact decreased with increasing concentrations of Myrrha. During the chemical examination of the Myrrha, it was noted that the material included 12 components known for their antimicrobial properties. The components described before are as follows: The specified chemical compounds consist of β-Elemene, β-bisabolene, Dihydro butyl bezodoxepin, Tetradecanol, methyl palmitate, Tribenzo-1,2,3,4,5,6 anthracene, 9-Eicosene, 2-bromo-4-fluoro-N-(thiophen-2-ylmethyl) aniline, Octadecenoic acid methyl ester, dehydroabietic acid, and Docosene.

**Conclusion:** The antimicrobial activity of five yeasts was assessed using the diffusion technique. The essential oil derived from two varieties of myrrha shown the most significant effects on *Candida tropicalis* (ATCC 66029), *Candida guilliermondii* (ATCC 6260), *Candida laurentii* (ATCC 18803), *C. neoformans* (ATCC 66031), and *Candida albicans* (ATCC 14053). Upon conducting a chemical examination of the myrrha, it was shown that it consists of 19 known components, of which 12 compounds have been proven by research to suppress the growth of microorganisms. Further research is needed to fully explore the potential of these water extracts as innovative antifungal medicines, including their effectiveness against various strains of *Candida* and the processes by which they work, as well as the primary components responsible for their action.

## Introduction

Myrrh resin, produced from a Bruseraceae plant, treats wounds, digestive difficulties, diarrhea, coughing, and chest diseases [1]. *Commiphora myrrh* treats wounds, oral ulcers, discomfort, fractures, stomach disorders, microbiological infections, and inflammation. Antiseptic, astringent, anthelmintic, carminative, emmenagogue, and expectorant is common medical terminology. The phytochemical examination found monoterpenoids, sesquiterpenoids, volatile/essential oil, diterpenoids, triterpenoids, and steroids. The essential oil is utilized in aromatherapy, perfumes, and cosmetics. The chemical is anti-inflammatory, antioxidant, anti-microbial, neuroprotective, anti-diabetic, anti-cancer, analgesic, and anti-parasitic. Recent studies demonstrate COVID-19 treatment works. This substance’s phytochemicals may be used to generate medical insecticides with anti-parasitic capabilities [2]. Batiha et al. [2] identified medication interactions. Sumerians used myrrh to treat tooth and intestinal worms about 1100 BC. Egyptians were myrrh embalmed.

In 1998, Stevenson [3] reported that the oil treats *Candida albicans*, Tinea pedis, and subcutaneous ulcers. The British Herbal Pharmacopoeia recommends myrrh tincture mouthwash for gingivitis and ulcers [4]. Blumenthal [5] reported in 1999 that the European Commission approved myrrh for minor pharyngeal and oral pain. Nomicos [6] states myrrh heals syphilis, leprosy, and rheumatism in Chinese medicine. In Somalia and Ethiopia, myrrh decoctions relieve stomachaches, de Rapper et al. [4] claim. Various studies show that terpenoids are anti-molluscic, hypoglycemic, local anesthetic, cytotoxic, and microbicidal [2].

Most healthy persons have cutaneous, oral, vaginal, and gastrointestinal *C. albicans*. In dysbiosis and immune suppression, symbiotic mode turns parasitic, producing invasive infections with significant morbidity and death from superficial to severe debilitation. *Candida albicans’* ability to transition between yeast, pseudohyphal, and hyphal forms is essential for host tissue invasion and parasitization, hyphae penetrate host tissues when yeast circulates [7].

Cryptococcus neoformans affects immunocompromised patients. Lethal meningoencephalitis results from CNS invasion [8].

The yeast-like environmental fungus *Papiliotrema laurentii* evolved from *Cryptococcus laurentii*. A restricted structure makes it non-pathogenic. Pigeon droppings contain this fungus [9,10]. Invasive *P. laurentii* infections affect immunocompromised persons. The first case of *P. laurentii* fungemia in a preterm infant with low birth weight in Kuwait and the Middle East was reported by [11]. Recent immunocompromised patients contracted invasive *P. laurentii* [12-16]. Additionally, *P. laurentii* may cause cutaneous infections [11, 17, 18].

Immunocompromised persons get skin infections from *Candida guilliermondii*. Human skin and mucosa contain saprophyte *M. guilliermondii*. Oncology, immunodeficiency, gastrointestinal or cardiovascular surgery, and chemotherapy may cause significant infections. The bulk of *M. guilliermondii* fatalities are cancer patients. *M. guilliermondii* infections have risen in recent decades. Low amphotericin B, fluconazole, micafungin, and anidulafungin response. Both preventive and empirical medicines increase *M. guilliermondii* MICs. Drug resistance worries researchers worldwide. Fixing this is their goal [19].

This product comprises essential oils, gum (30-60% water solubility), myrrhol (3-8% ether), and resin, myrrhine. Burseraceae species *Commiphora* produce fragrant, transuding resins from stem bark. Yellow-to-reddish-brown resin particles. Bitter taste and pungent balsamic aroma are pleasant [2,20]. Heerabolene, elemol, acadinene, cuminaldehyde, eugenol, and furano-sesquiterpenes such furanodienone, furanodiene, curzerenone, and lindestrene are in *Commiphora myrrh* resin [21] the volatile oils are mono- and sesquiterpenoids. Gas chromatography finds monoterpenoids. Many gas chromatography experiments have explored *Commiphora* species’ volatile oils. *C. myrrha*, quadricincta, holtziana, guidottii, kataf, sphaerocarpa investigated. Mono-terpenoids found in literature include camphene, myrcene, limonene, α-pinene, and β-pinene. *Commiphora* volatile oils vary widely. Volatile oil demands low oxidation sesquiterpenoids. β - selinene, β-Elemene, α-humulene, α-copaene, and germacrene B are common in *Commiphora* species’ volatile oils. Furanosesquiterpenoids characterize *Commiphora*.

Biofilm-related *Candida albicans* disorders are difficult to treat owing to antifungal drug resistance. The search for innovative phytotherapeutic approaches employing medicinal plants and essential oils has begun [22]. Rahman et al. [23] identified four antimicrobial *Commiphora* mol terpenes: mansumbinone, 3,4-seco-mansumbinoic acid, β-elemene and the latter is antibacterial. It slightly increases ciprofloxacin and tetracycline against *Salmonella* strains SL1344 and L10 and eight times Norfloxacin against *Staphylococcus* [23]. Germano et al. [24] found *C. myrrh* extract and essential oil expectorants treat chest infections. Su et al. [25] found myrrh aromatic gum resin causes chest infections. The resin fights chest infection-causing germs and fungus by inhibiting inflammation and cytotoxicity. *C. myrrh* resin and extract decreased inflammation and chest discomfort [26]. Sore throat and chest infection myrrh. Decreases inflammation [27]. In diffusion trials, pe myrrh inhibited *C. albicans*. Water extracts stopped strains. Myrrh is more effective against *C. albicans* (9 mm zone of inhibition, 20 mg/mL) for gingivitis, pharyngitis, phyorrhoea, and sinusitis [3] *C. albicans* antifungal activity matches previous observations [28] of *Commiphora. Commiphora* resin and Fiorano sesquiterpenoids have antimicrobial and antifungal properties. This study characterized the chemical constituents of the aqueous extract of myrrh resin and demonstrated its antibiotic potential by inhibiting many pathogenic yeasts.

Based on this premise, this research suggests doing more investigations to separate and isolate the natural active components from the plant extract. These components help save lives and aid in discovering alternatives to combat the emergence and spread of antibiotic-resistant types of microbes.

## MATERIALS AND METHODS

### Plant samples

The plant samples of *Commiphora myrrha* and *Commiphora molmol* came from stores in Dammam city. We kept the items we bought at 4°C until we could test them.

### Preparing the extracts

#### Aqueous extract

*Commiphora myrrha* and *C. molmol* oleo-gum resins Dammam supplied additional samples for this study. Myrrha powder 1, 2, and 5g was dissolved in 10 mL sterilized distilled water. They were gauze-filtered after 24 hours of ambient soaking [30].

##### Fungal strains

Five fungal strains were obtained from School of Medicine, King Saud University, Riyadh city. These fungi were different strains of *Candida albicans* (ATCC 14053), *C. neoformans* (ATCC 66031), *Candida laurentii* (ATCC 18803), *Candida guilliermondii* (ATCC 6260), and *Candida tropicalis* (ATCC 66029).

Each strain was subculture on Sabouraud’s dextrose agar (SDA) (Scharlau, Spain) medium and incubated at 37°C for 24 h to obtain inoculums for testing [31].

### Testing the effectiveness of aqueous extracts

Well-diffusion agar was used [31], after adding 1 ml of the fungal suspension of each Candida individually to a sterile plastic petri dish, 10 ml of the medium was poured off and left to solidify. Using a sterile corkborer, a hole of half a centimeter in diameter was made in the agar, and the myrrh extract was placed inside it and incubated for 48 hours. The plates were incubated at a temperature of 37°C. The area occupied was calculated to record the results.

### Detecting the composition of the chemical materials of Myrrha resin aqueous extract

Mass spectrum was used to analyze myrrha’s chemical composition for antimicrobial chemicals. Analyzing the sample’s mass spectrum using GC-SM assessed its chemical composition. After identifying the samples’ primary ingredient, the inhibitive chemicals are determined [32].

## STATISTICAL ANALYSIS

Statistics were done using a properly randomized technique and three treatment duplicates. L.S.D. in SPSS 16 was used to compare data at 0.05 probability according to Norusis [33].

## RESULTS AND DISCUSSION

### Effect of tested Myrrha aqueous extracts

The impact of tested Myrrha aqueous extracts on five isolates was examined using the agar well diffusion technique, chosen for its reliability, simplicity, and clear outcomes. The findings were established by measuring the area of the inhibitory zone after a period of 48 hours. The findings shown in Table (1) indicate that an inhibitory zone was identified for both kinds of bitter substances against the tested candida strains. Furthermore, the inhibitory impact was found to be directly proportional to the dose utilized, ranging from 10% to 30%.

**Table-1:**
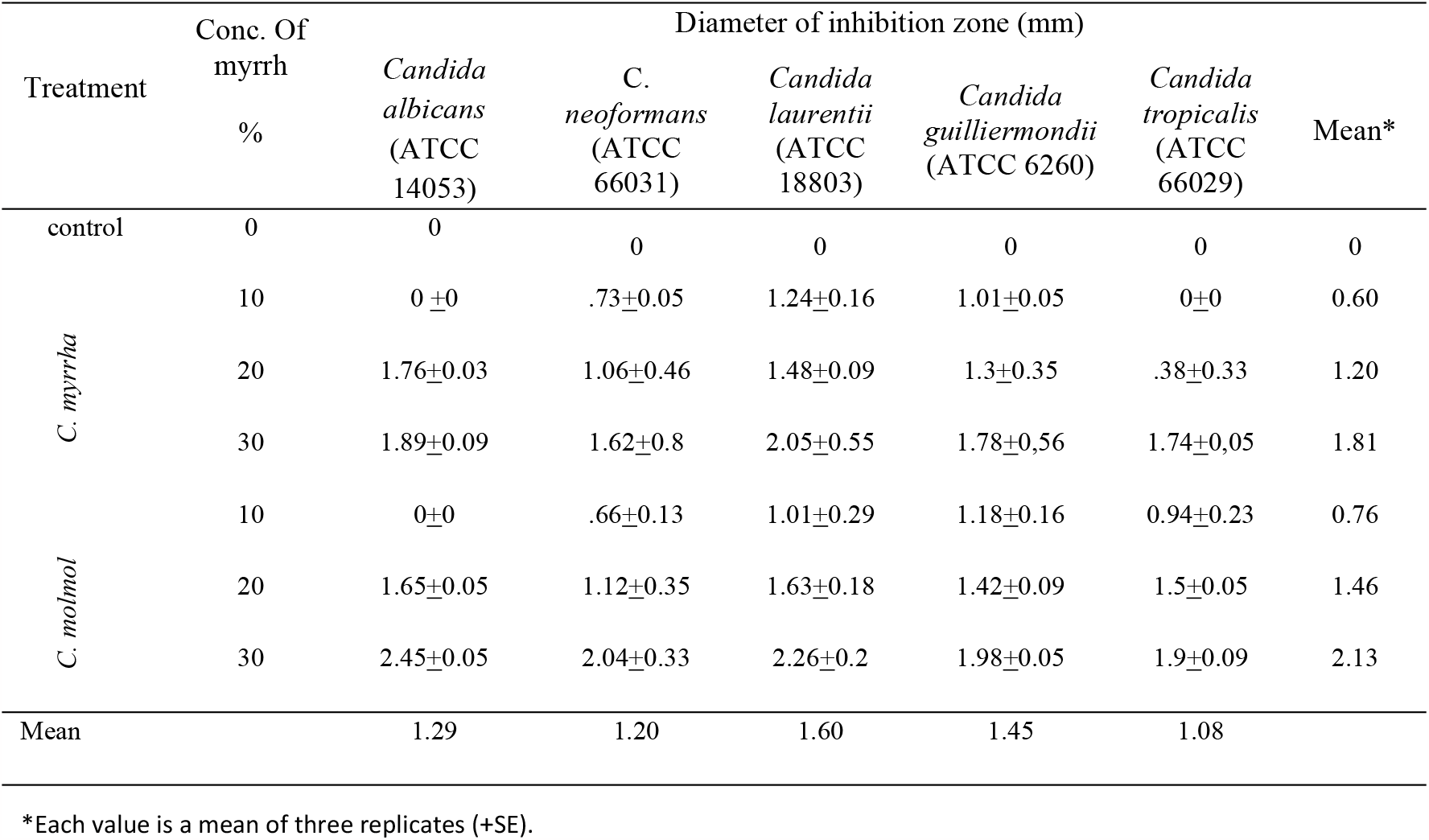
Inhibition zone (mm) of Myrrh extracts at various concentration on some Candidiasis.

The inhibitory zone diameters for *Candida albicans* (ATCC 14053), *C. neoformans* (ATCC 66031), *Candida laurentii* (ATCC 18803), *Candida guilliermondii* (ATCC 6260), and *Candida tropicalis* (ATCC 66029) were 2.45 mm, 2.04 mm, 2.26 mm, 1.98 mm, and 1.9 mm, respectively, at a 30% concentration of *C. molmol*. However, the use of C. myrrha aqueous extract resulted in a reduction in the zone of inhibition to 1.89, 1.62, 2.05, 1.78, and 1.74, respectively. Akintobi et al [34] have achieved comparable findings, indicating that both forms of Myrrha have had an impact on *C. albicans*. The area of inhibition measures 1.76 and 2.01 cm2 for doses of 250 and 1000 mg/mL respectively for the first kind of Myrrha. For the second type, the area of inhibition measures 1.53 and 1.76 cm^2^ consecutively.

The results obtained by evaluating the perception of candida in reaction to these extracts align with the conclusions of other studies, including as [34-37]. The beneficial effects of *Candida* may be ascribed to the existence of microbiologically resistant chemicals in these extracts, such as volatile oils, terpenes, phenols, flavonoids, and saponins [38].

The pathogenicity of Candida species is due to various virulence factors, including their ability to evade the immune system, adhere to surfaces, promote the growth of hyphae, form biofilms on medical devices and host tissue, and produce hydrolytic enzymes such as proteases, phospholipases, and haemolysin, which can cause tissue damage [39]. persons with weakened immune systems or diabetes mellitus are more vulnerable to getting mucosal or systemic infections compared to healthy persons [40]. Non-albicans species are contributing to a concerning increase in infections. *Candida glabrata* is the second most common species of candidiasis or vaginal candidiasis, behind *C. tropicalis*, which is the third most widespread species, and *C. albicans. Candida parapsilosis* has become the dominant fungal infection in newborns and infants in certain medical institutions [41].

### Chemical composition of myrrha resin in the aqueous extract

The obtained results demonstrated that the aqueous extract of both types of myrrha contains substances that hinder the proliferation of the five types of *candida*. In order to determine the nature of these compounds, a chemical analysis was conducted on the composition of the myrrha using the mass spectrum method (figure 1). The presence of 12 compounds with known antimicrobial properties has been demonstrated. These compounds are: β-Elemene, β-bisabolene, Dihydro butyl bezodoxepin, Tetradecanol, methyl palmitate, Tribenzo-1,2,3,4,5,6 anthracene, 9-Eicosene, 2-bromo-4-fluoro-N-(thiophen-2-ylmethyl) aniline, Octadecenoic acid methyl ester, dehydroabietic acid, and Docosene. The extract’s inhibitory activity may be attributed to the presence of abundant terpene single molecules [42]. These chemicals possess the capacity to inhibit many forms of yeast, which is known as their ability to examine the cell wall. This also results in the attenuation of cellular biological processes by interfering with the cytoplasmic membrane’s protein synthesis process, so slowing and halting the process. This also impedes the active transport of ions and salts via the membrane [43].

**Figure. 1.**
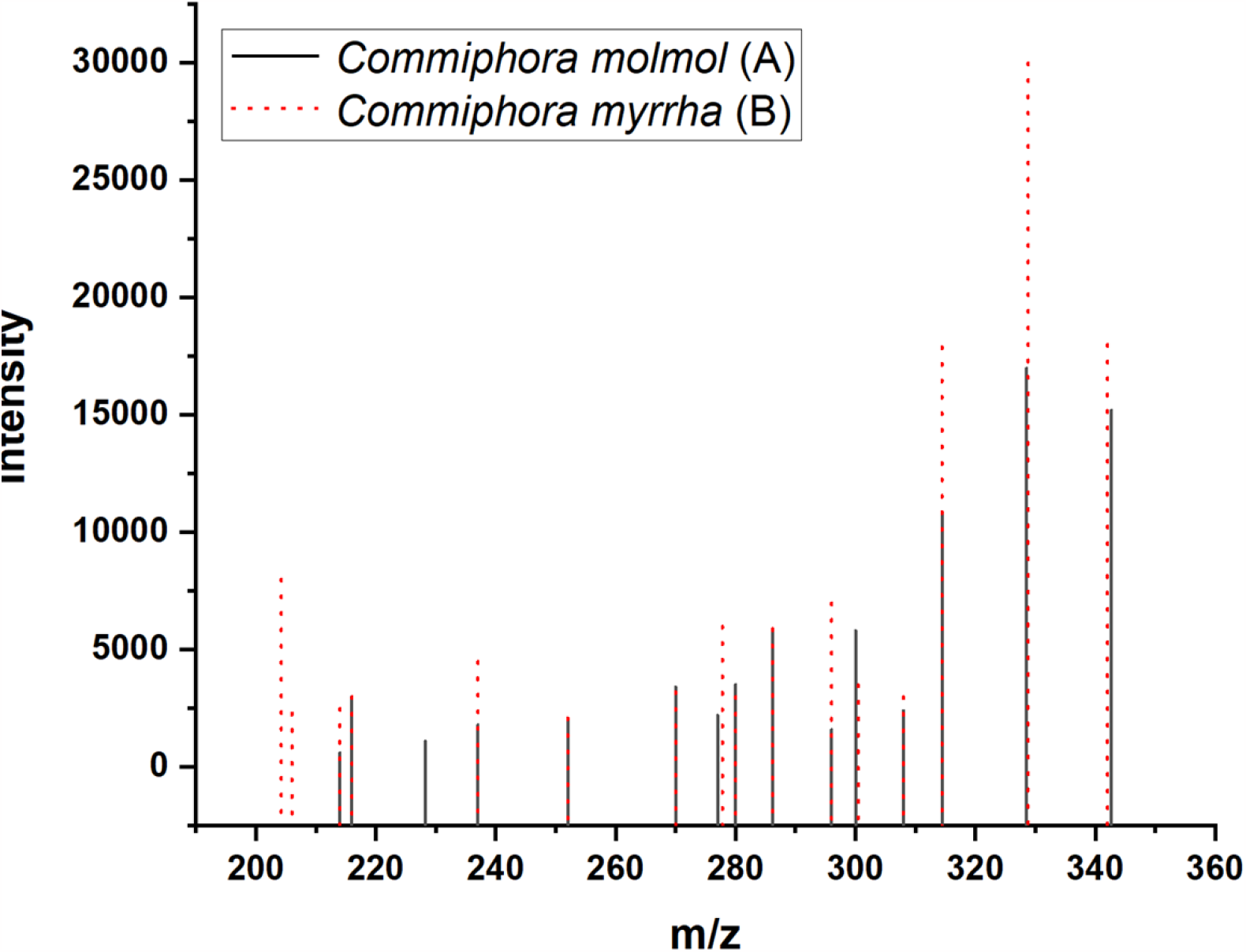
Chromatograms differentiate the primary constituent of *C. myrrha* and *C. molmol* via mass spectrometry.

The specific strategies used by microbes to survive the effects of microbial antibiotics are still unclear and subject to debate [35]. Conversely, the chemical constituents of plants serve to safeguard them from internal microbial assaults. Nevertheless, some components within these substances possess inherent significance as natural chemical compounds that serve to protect the human body against microbial invasions [44].

The results obtained from the feeling of candida of these extracts are consistent with the findings of other studies [34-36]. The favorable findings of *Candida* may be attributed to the presence of potent components in these extracts that are resistant to microbiological agents, such as volatile oils, terpenes, phenols, flavonoids, and saponins [38].

Figure (1) shows the GC-MS chromatographic analysis of the ethanolic extract of two varieties of Mrayh. The analysis was conducted using a GC-MS equipment to first identify and describe its components. Table (2) presented the constituents of the raw Mayrah ethanolic extract, revealing the identification of 19 subcomponents with molecular weights ranging from 146 to 410.

**Table (2):**
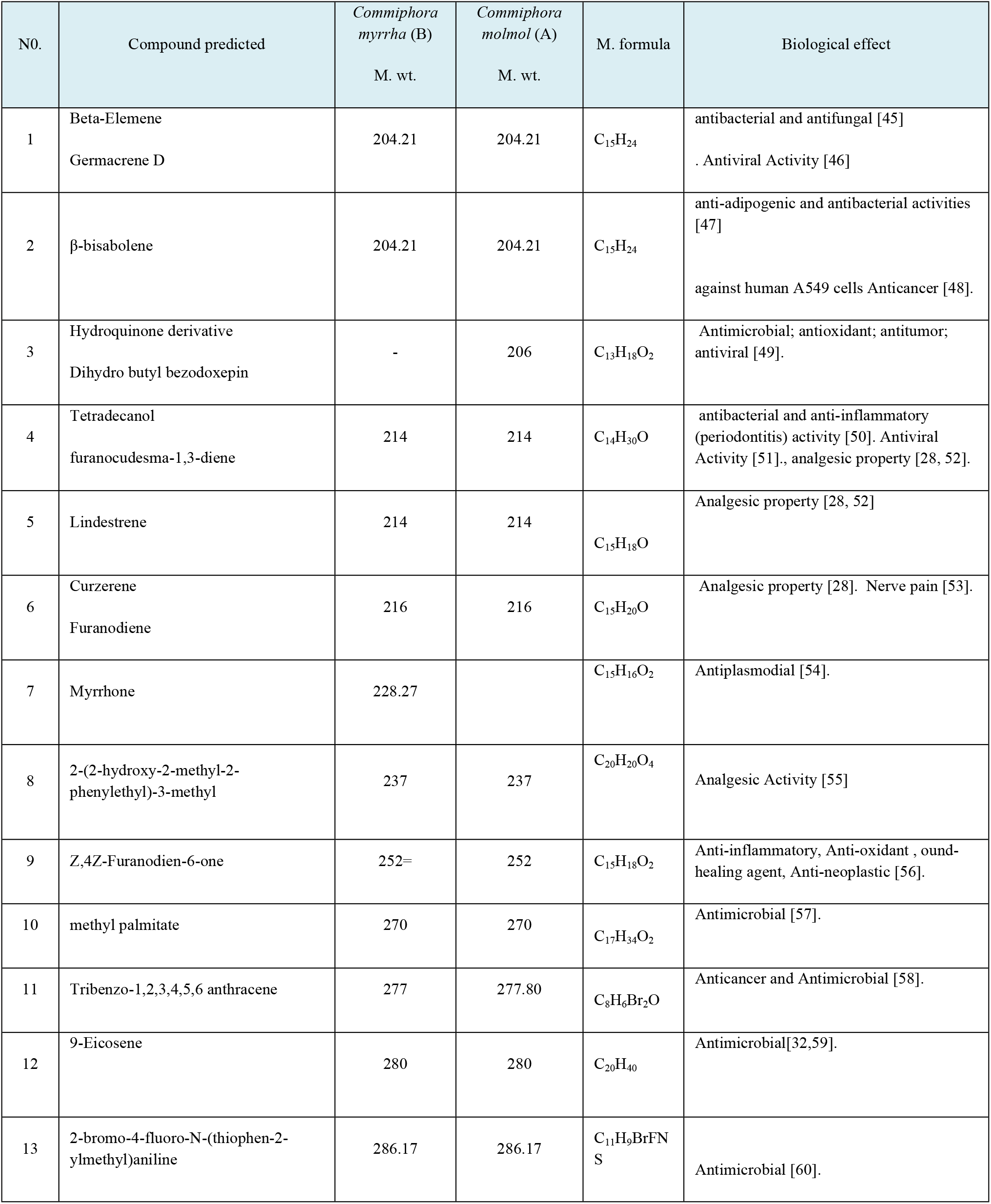
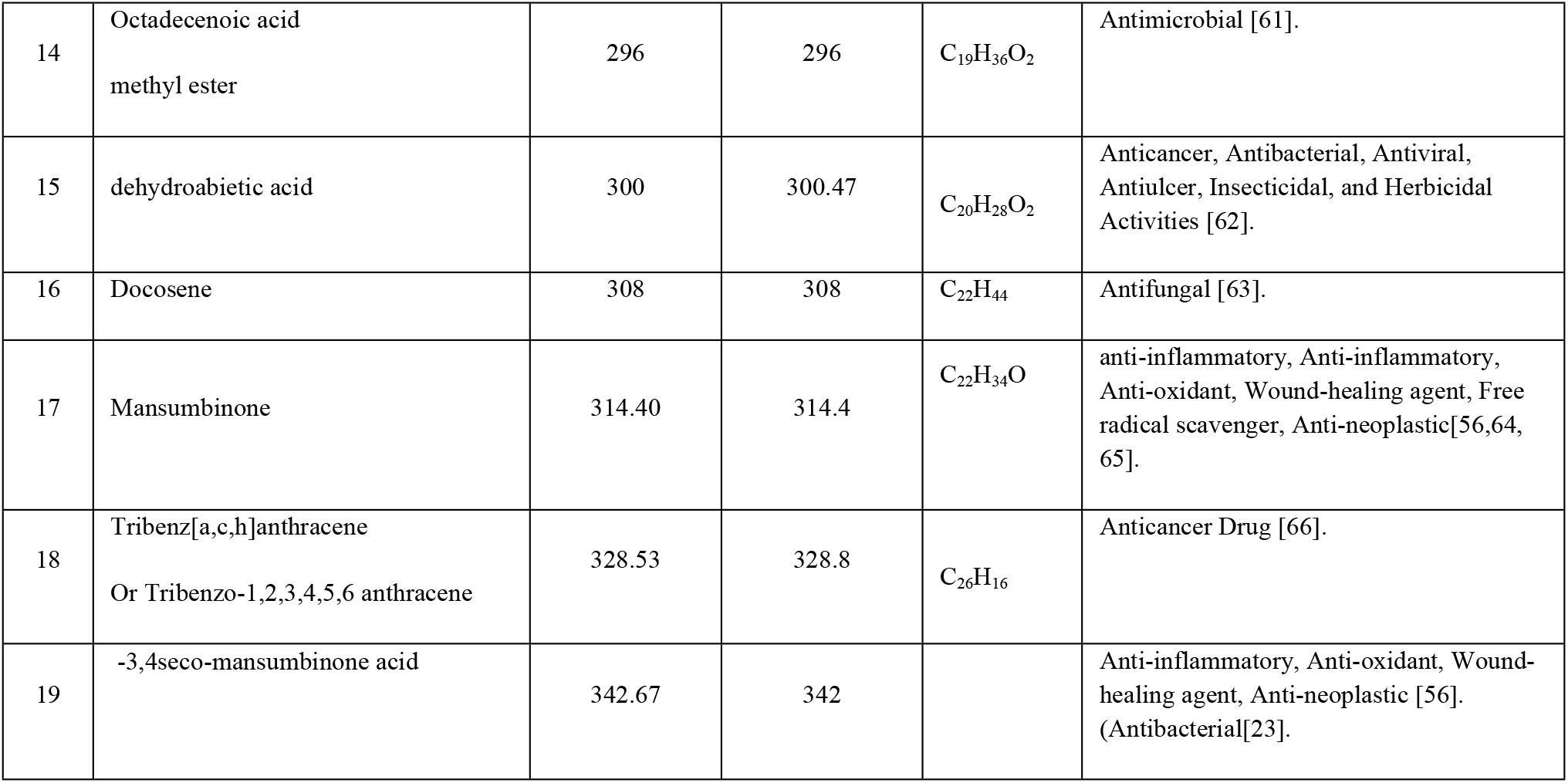
Analyzing the GC-MS chromatograph reveals the expected subcomponents of the SFE *C. myrrha* and *C. molmol* extract.

These subcomponents include fatty acids and their precursors, with some of them being categorized as polyunsaturated fatty acids (PUFAs), such as Linoleic acid. Eugenol and trans isoeugenol are phenolic chemicals, whereas squalene is an alkene categorized as an isoprenoid substance.

Table 2 and Figure 1, 2 show the chemical composition of the myrrh plant extract. The GC-MS study revealed the presence of 19 compounds, including 12 compounds with antifungal properties. All of these chemicals are classified as fatty acids and their precursors, as well as phenolics and flavonoids.

**Figure 2:**
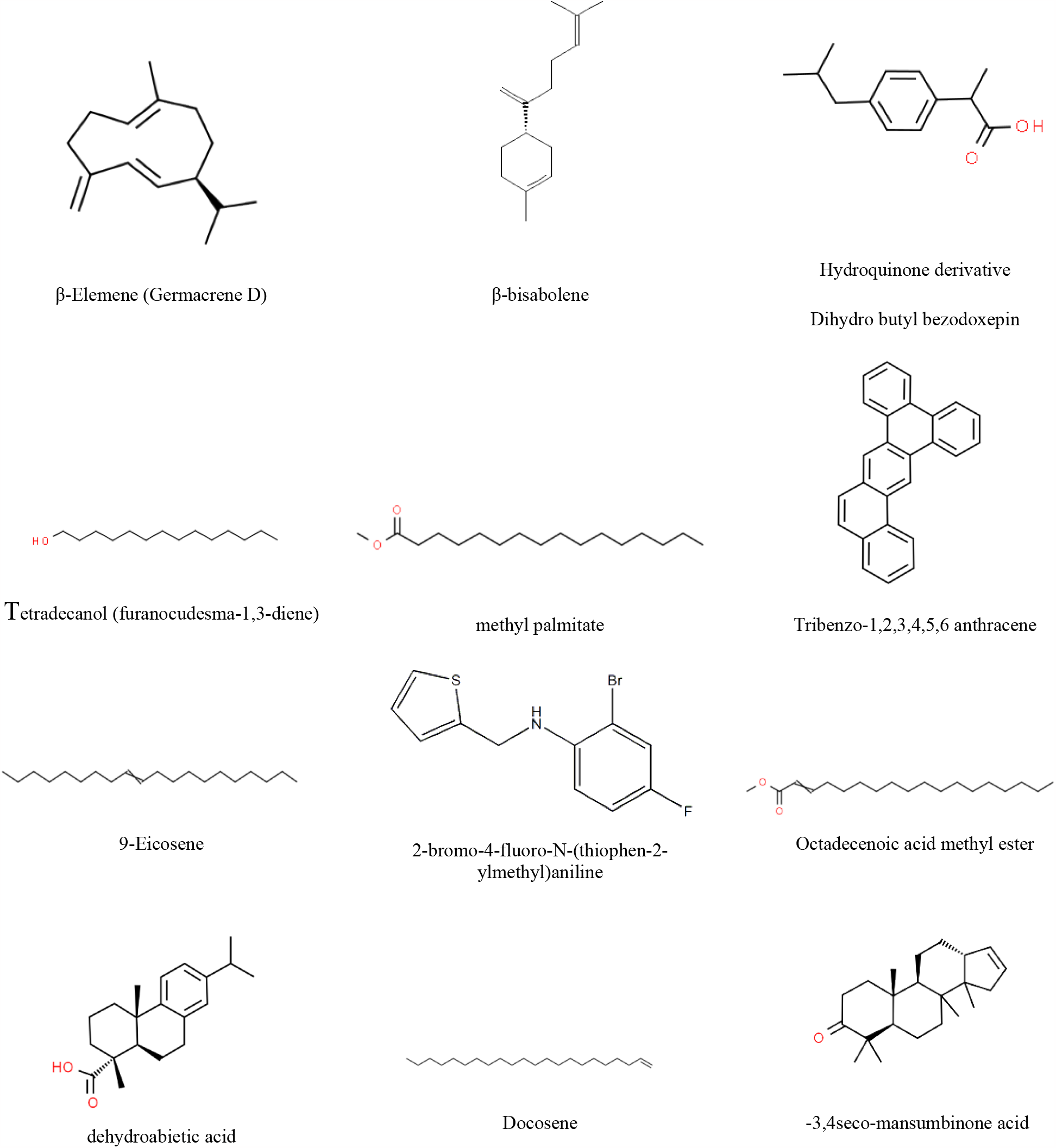
The Chemical constructions of biologically active compounds extracted from *Commiphora myrrha* and *C. molmol* oleo-gum resins.

The aqueous extracts of *Commiphora myrrha* and *Commiphora molol* reduced the activity of the tested fungus significantly. This might be due to fungal cell wall damage or even damage to internal organelles such as mitochondria, which hampered their ability to gather energy for development. Murakami et al.,[67] noted that the essential oil derived from *C. myrrha* led to substantial decreases in the activity of mitochondrial dehydrogenase in *C. albicans*, hence affecting its potential to produce energy. Furthermore, a considerable reduction in biofilm production was noted, with a 62% reduction. There has been no prior research on the effect of *C. myrrha* essential oil on *C. albicans* that we are aware of. Furanoeudesma-1, 3-diene (17.65%), curzerene (12.97%), β-elemene (12.70%), and germacrene B, D, and A (12.15%, 9.13%, and 5.87%, respectively) are the primary components of *C. myrrha* essential oil. Antifungal effects of curzerene and β-elemene have been shown. Nonetheless, there has been little research on the effect of the compounds found in *C. myrrha* essential oil on *Candida albicans*. As a result, additional research is required to explore into this area [67, 68, 69].

## Conclusion

Plant resin is a valuable botanical resource in traditional medicine. *Commiphora* is anti-inflammatory, antioxidant, anti-microbial, neuroprotective, anti-diabetic, anti-cancer, analgesic, and anti-parasitic. Additionally, it may fight respiratory infections like COVID-19. These pharmacological effects are caused by terpenoids (monoterpenoids, sesquiterpenoids, volatile/essential oils), diterpenoids, triterpenoids, and steroids. The essential oil is used in aromatherapy, scent, and cosmetics. Pharmaceuticals’ rich phytochemical components might be used as insecticides as medicine research advances, due to their great anti-parasitic effectiveness and ability to grasp medication interactions fully. Five yeasts were tested for antifungal activity utilizing diffusion. The essential oil from two myrrha types had the greatest impact on *Candida tropicalis* (ATCC 66029), *Candida guilliermondii* (ATCC 6260), *Candida laurentii* (ATCC 18803), *C. neoformans* (ATCC 66031), and *Candida albicans*. A chemical analysis of myrrha revealed 19 components, 12 of which have been found to limit microbe development. Further study is required to thoroughly examine these water extracts’ potential as revolutionary anti-fungal drugs, including their efficiency against diverse Candida strains, their mechanisms of action, and their main components.

## Acknowledgments

The author is thankful to Department of Pathology, School of Medicine, King Saud University, Riyadh 11461, Saudi Arabia for providing the cultures of *Candida* strains.

## Data Availability

Data used to support the findings of this study are included within the article.

## Conflicts of Interest

The author declares that there are no conflicts of interest.

## Notes

### Competing Interest Statement

The authors have declared no competing interest.

